# Neotenic transcriptomic features in the adult turquoise killifish brain

**DOI:** 10.1101/2025.03.24.645095

**Authors:** Ana María Burgos-Ruiz, Silvia Naranjo, Maria del Valle Hernández-Pérez, Lorena Ardila-García, Marta Moreno-Oñate, Ana Fernández-Miñán, Alejandro de los Reyes-Benítez, Javier López-Ríos, Gretel Nusspaumer, Juan J. Tena

## Abstract

Aging is one of the main challenges facing modern society. Understanding the cellular processes that occur in the later stages of life is essential to address age-related diseases. Although aging is a global process that affects the entire body, the brain is one of the most sensitive organs and is affected by numerous degenerative diseases. In this study, we investigate cell types and genes that are particularly sensitive to the aging process. To this end, we performed a time-course single-cell RNAseq experiment series in the brain of the killifish, the vertebrate model with the shortest lifespan, which makes it an excellent model for aging studies. Our analysis reveals that non-glial progenitor cells are among those populations that change the most between young and old animals. Furthermore, we identify specialized stromal clusters that seem to support a primitive hematopoietic program in the brain, which is active only during the embryonic stages in other vertebrate species. Our results show that the expression of embryonic genes in the adult brain appears to be a general feature in killifish, and that several cell populations in the adult killifish brain show a higher level of similarity to the zebrafish embryonic populations than to adult ones. Our study suggests that adult killifish maintain a neotenic gene expression status in the brain that may help in sustaining their characteristic high proliferation and metabolic rates, as well as combat the detrimental effects of this high metabolism on cells, especially at advanced ages.

## Introduction

Aging is considered one of the most important challenges from the social, economic and medical point of view. The aging process is characterized by a progressive functional decline across tissues and organs, as a consequence of the general deterioration of the physiology of cells. This leads to the development of numerous pathologies such as neurodegenerative and cardiovascular diseases, cancer and diabetes. Although aging is considered a global process that affects all cells in the body, different organs and tissues show different sensitivity to this process (López-Otín et al. 2013). Among them, the brain is particularly sensitive to age (Mattson and Magnus 2006; Morrison and Baxter 2012), and is associated with many age-related diseases, such as Parkinson, Alzheimer and other neurodegenerative diseases.

Different animal models have been used for the study of the aging process and related diseases. Traditional vertebrate models such as zebrafish or mouse have relatively long lives, around 5.5 and 4 years, respectively (Hu and Brunet 2018), what considerably complicates aging experimentation in laboratory conditions. This has resulted in aging studies in vertebrates being fairly limited so far, in particular at advanced or geriatric ages, and mainly focused on confirming results obtained in yeast or invertebrate models, like worms or flies, which present much shorter lifespans (Taormina et al. 2019).

Recently, the turquoise killifish *Nothobranchius furzeri* has stood out as a new model with great potential for the study of longevity in vertebrates (Valdesalici and Cellerino 2003; Reichwald et al. 2015; Harel, Valenzano, and Brunet 2016). The killifish, recognized as the shortest-lived vertebrate capable of reproducing in captivity and with a lifespan ranging from 3 to 8 months (Hu and Brunet 2018), undergoes a rapid natural aging process as an adaptive response to its environment. Despite their short lifespan compared to other vertebrates, killifish show many of the physiological aging traits characteristic of other vertebrates (Hu and Brunet 2018).

The killifish lives in ephemeral ponds created by seasonal rains, which dry up within months, restricting survival and reproduction to a narrow timeframe. Only species capable of giving rise to a new generation within such a short-lived habitat can successfully establish themselves there, making rapid sexual maturation crucial. The turquoise killifish is the fastest-maturing vertebrate, reaching sexual maturity only 14 days post-hatching (Hu and Brunet 2018). The rapid maturation of these animals is driven by an explosive growth rate. This accelerated development is also reflected in the brain, as studies have shown that the dorsal telencephalon (pallium) grows three times faster than in zebrafish (Coolen et al. 2020).

Whether the rapid aging observed in killifish is a consequence of their explosive growth, or if these processes occur independently, is not known yet. It is also unknown if rapid aging preferentially affects specific cell populations or signaling/metabolic pathways in the killifish brain. To gain insight into these open questions, we conducted an in-depth study of gene expression and cellular composition in the killifish brain as aging progresses. To this end, we resorted to single-cell RNA sequencing (scRNA-seq) to analyze transcriptomic variations across different adult life stages, from youth to senescence, and to examine changes in the size and distribution of specific cell populations. Our study reveals a dynamic scenario as the killifish brain ages, and uncovers a set of neotenic features at both the gene expression and cell population levels.

## Results

### Synexpression groups of genes in the killifish adult forebrain

To identify global trends in the transcriptional programs operating on the different cell populations of the brain along the lifespan of killifish, we first performed bulk RNAseq assays with forebrain samples at four time points (1.5, 2.5, 4 and 5 months), since the animals have reached the sexual maturity, until clear signs of senescence, like loss of pigment and muscular tissue or reduced activity, were observed, and the animals are considered geriatrics. After clustering all genes depending on their transcriptional behavior through life, we identified several gene synexpression groups with similar transcriptional dynamics (Supp. Fig. 1 and Supp. Table 1).

Several interesting groups of genes were identified in this analysis. For example, the group 15, with a marked increasing expression profile during the animal lifetime, clearly reflects functions related to hematopoiesis, as revealed by gene ontology analysis (group 15, Fig. 1A, Supp. Fig. 1, Supp. Table 1). Group 2, which presented high expression in young animals and lower in later stages, although with an increasing tendency in old animals, was related to early erythropoiesis (Fig. 1B, Supp. Fig. 1, Supp. Table 1). Other interesting synexpression groups identified were: group 16, which showed high expression levels in the two first time points and lower in old animals, and was enriched in GO terms related to cell division, which may be attributed to a reduction in the brain’s growth rate; group 14 was composed by genes highly expressed at the first and the last timepoints, and was enriched in terms related to protein synthesis, which could be linked to initial growth and later adaptive responses to stress; and group 1, which contains genes with much higher expression levels in the first time point than in the rest, and was enriched in GO terms related to early neural development, which would be expected to be active at earlier embryonic stages rather than in already sexually mature individuals (Fig. 1C, Supp. Fig 1, Supp. Table 1).

**Figure 1.**
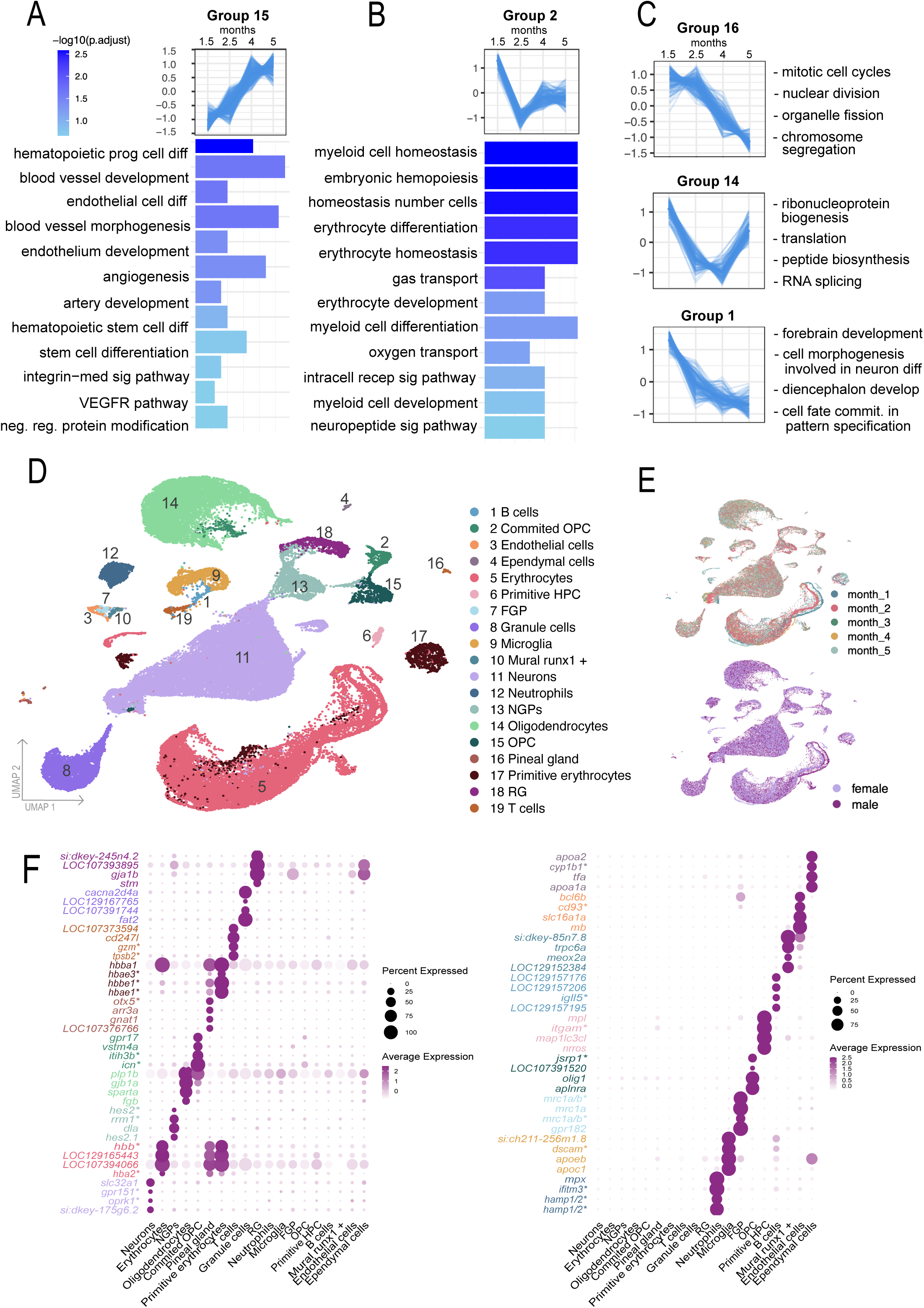
Killifsh brain RNAseq and scRNAseq analyses. **(A, B)** Differentially expressed genes (DEGs) were identified through RNAseq analysis of four time points, comparing each time point to the baseline. Temporal expression dynamics were grouped into 20 clusters using k-means clustering. The average expression trend of genes in synexpression groups 15 **(A)** and 2 **(B)** is presented, along with the top 12 associated Gene Ontology (GO) terms (Biological Process). **(C)** Temporal trends of three additional gene groups (16, 14, and 1) and some representative associated GO terms (Biological Process) from their top 12 annotations. **(D)** Annotated UMAP of integrated data from seven samples, highlighting the identified cell types present across all samples. **(E)** UMAP showing the distribution of cells according to the age and the sex of the sample. **(F)** Dot plot summarizing the expression of key marker genes across the major cell types identified in the killifish brain. The size of each dot represents the proportion of cells within a given cell type expressing the marker gene, while the color intensity reflects the average expression level. Gene names marked with an asterisk come from their orthologs from zebrafish.

### Single-cell resolution transcriptomic profiling throughout brain ageing

To further investigate which cell populations within the brain were the most sensitive to the aging process, and which genes participate in the main changes we observed in these cells, with single-cell resolution, we performed single-cell RNAseq (scRNAseq) profiling of whole brains, all along the adult life of killifish, from 1 to 5 months. Furthermore, since sex is an important factor to take into account in aging-related diseases (Regitz-Zagrosek 2012; Braunwald 2013; Garate-Carrillo et al. 2020), we used male and female samples. In total, 7 single-cell experiments were performed: 5 from males (months 1, 2, 3, 4, and 5) and 2 from females (months 2 and 5) (Supp. Table 2). After sequencing and quality control, a total of 59,985 cells were obtained, distributed across the seven samples (Fig. 1D, E, Supp. Fig. 2A, B).

Although killifish genome has been previously sequenced and annotated, generating a high-quality, very complete assembly (https://www.ncbi.nlm.nih.gov/datasets/genome/GCF_027789165.1/), Gene Ontology (GO) terms libraries are still missing for this species, which makes cluster annotation for scRNAseq studies challenging. Based on already known markers for different cell types in the brain and comparing them with the most prominent differential genes from each cluster, we were able to annotate all the 19 clusters identified in our analyses (Fig. 1D and Supp. Fig. 2C). Among them, we could pinpoint some clusters from the neuronal lineage, such as mature neurons, non glial progenitors (NGP), granule cells and pineal gland cells (which might have a mixed origin (Staudt et al. 2019)); several clusters from the glial lineage, like oligodendrocytes and oligodendrocyte precursor cells (OPC), microglia and radial glia (RG), among others; and clusters from hematopoietic lineage, from both the lymphoid branch, like B and T cells, and the myeloid one, such as erythrocytes and neutrophils.

A complete analysis of differentially expressed genes (DEG) revealed groups of marker genes specifically expressed in each cell type (Fig. 1F, Supp. Fig. 2D and Supp. Table 3), which constitute a valuable resource for future research. For example, to better characterize the NGP cells in killifish—remarkably distinct and not yet identified in adults of other species— our analysis shows that their top three marker genes are *hes2*, *rrm1*, and *dla*. *hes2* acts as a transcriptional repressor regulating neurogenesis and cell fate, suggesting a developmental role; *rrm1*, essential for DNA synthesis and repair, indicates proliferative capacity, while *dla*, a Notch pathway component, highlights involvement in cell differentiation. Most importantly, we found specialized angiogenic-related (cluster 3, 10) as well as some hematopoietic clusters (6, 7 and 17), which aligns well with the identification of synexpression groups enriched in blood vessel and hematopoiesis associated GO terms identified through bulk RNAseq analysis (Fig. 1A, B).

After cluster annotation, our first analysis compared the number of cells in each of the clusters throughout different timepoints and between males and females (Fig. 1E, Fig. 2A, B, Supp. Fig. 3A). Several clusters showed statistically-significant differences in the cell number at two months compared to 5 months, indicating that these cells are undergoing changes in population size during aging. This is the case of primitive erythrocytes and fluorescent granular perithelial cells (FGP) that experience a decrease in cell number during aging in males, or granule cells and primitive hematopoietic progenitor cells (primitive HPC) in females, which increase in cell number during adult life (Fig. 2B).

**Figure 2.**
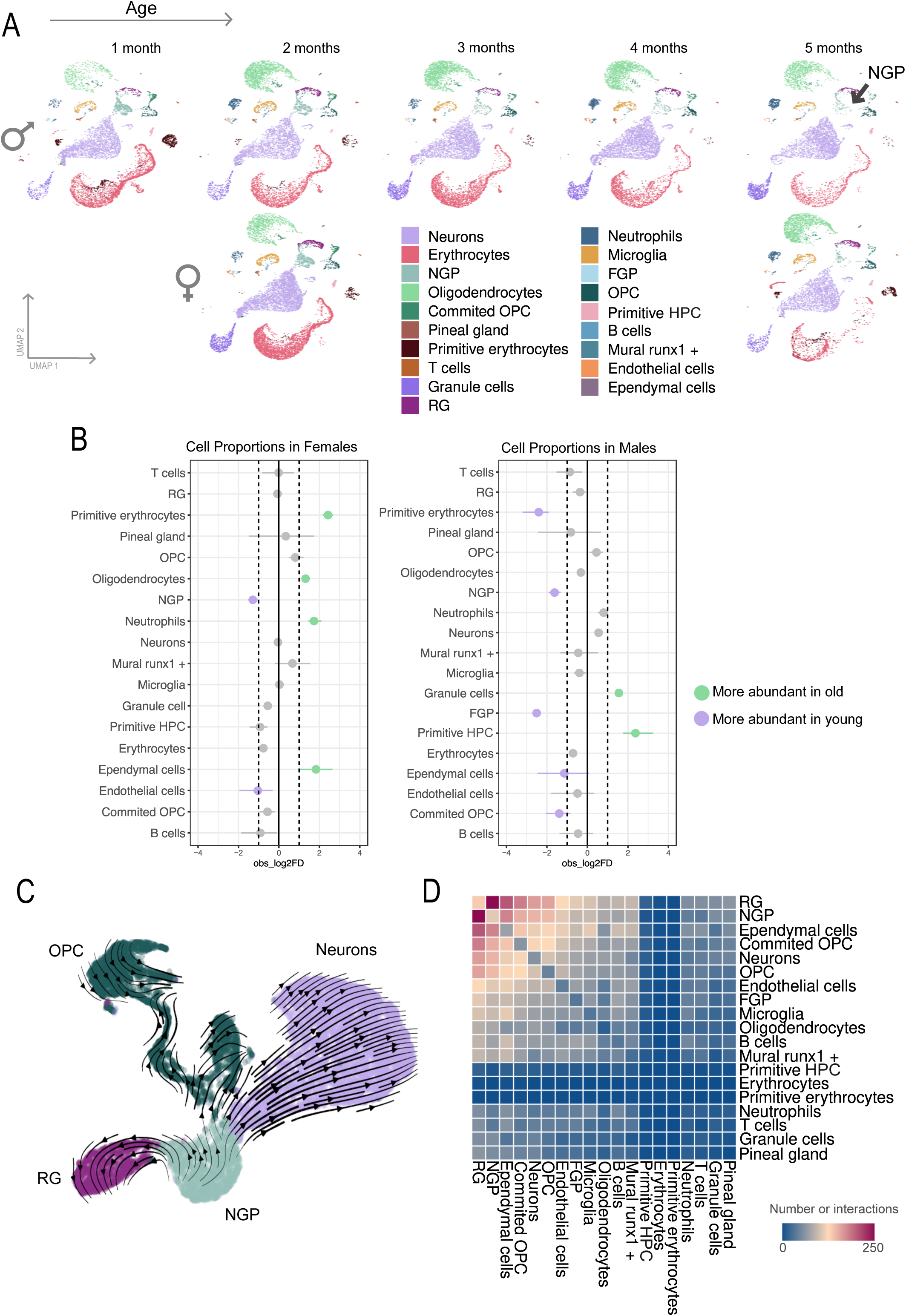
Cell population dynamics in scRNAseq clusters. **(A)** UMAPs split by age and sex, illustrating changes in the dynamics of cell population sizes across samples. **(B)** Differences in cell type proportions between ages for each sex. Left-shifted cell types are more abundant in young fish, whereas right-shifted cell types are more abundant in old fish. Dashed lines represent significance thresholds of a 1.5-fold change in cell abundance at p < 0.05. **(C)** Representation of cellular trajectories among neural progenitor cells (NGP), oligodendrocyte precursor cells (OPC), radial glia (RG), and neurons. **(D)** Number of interactions between cell types in all samples.

Only NGP decreased significantly their numbers in both males and females during aging, although they did not disappear even at the last time point, which is interesting because this population represents the capacity of killifish of neuronal renewal during their entire life. To further investigate the NGP population, we performed trajectory analyses. Cell-fate analysis with scVelo software (Bergen et al. 2020) showed that NGP give rise to neurons and radial glia (RG), which was previously reported by other authors (Bergen et al. 2020). However, there is a third population of cells that, according to these predictions, is derived from NGP, the oligodendrocyte precursor cells (OPC) (Fig. 2C), which might originate from different sources during embryogenesis in other vertebrate species (Beiter et al. 2022; Johansson et al. 1999; Dawson et al. 2003). NGP also exhibit extensive cell-cell predicted interactions with RG, and together they represent the two cell types with the highest number of interactions in our dataset, as revealed by our analysis using CellPhoneDB software (Vento-Tormo et al. 2018) (Fig. 2D). RG cells appear to interact with NGP and regulate their differentiation to other cell types and/or their precursor state maintenance through Wnt/Fzd pathway-mediated interactions (Supp. Fig. 2B).

Next, we analyzed changes in gene expression across different clusters throughout lifetime in both males and females. For this, we first used the R package Augur (Skinnider et al. 2021), a machine-learning-based tool that ranks cell types according to their sensitivity to a given biological perturbation—in this case, aging. Using this approach, we could identify NGP, primitive erythrocytes and B cells as the most sensitive cell types to the aging process in killifish (Fig. 3A). Indeed, NGP were the cells with the highest number of differential genes between young and old animals (Fig. 3B), as calculated using pseudobulk-RNAseq with the muscat software (Helena L., Charlotte Soneson, Pierre-Luc Germain, Mark D. 2019). For example, *chek1,* which plays a crucial role in DNA damage response and cell cycle checkpoint control, decreased its expression in NGP and OPC as animals got older. s*carb2a*, important for lysosomal function and autophagy, was found to be downregulated during aging in NGP, neurons, OPC and RG clusters, while *antxr2a* (involved in cell adhesion and extracellular matrix interactions) and *pde7b* (implicated in cellular responses to stress and inflammation) were upregulated in neurons (Fig. 3C). This type of analysis highlights how different cell types undergo distinct changes in gene expression during aging, as illustrated by the top differentially expressed genes (DEGs) for each cell type (Supp. Fig. 3B). These results were validated for some of the DEG by qPCRs in males and females at 1.5 and 5 months (Supp. Fig. 3A). Most of the tested genes showed in the qPCRs analysis the same trend as in the scRNAseq, but we also found two cases in which the gene behaved differently in male and female samples (*Ppde7b*, *Hs70prot1lk,* Fig. 3C, Supp. Fig. 3A). This could be explained because the expression levels of these genes change very quickly after the first month, and the samples for performing qPCRs may have been collected just after this inflection point, reflecting an increasing trend in males and the opposite in females.

**Figure 3.**
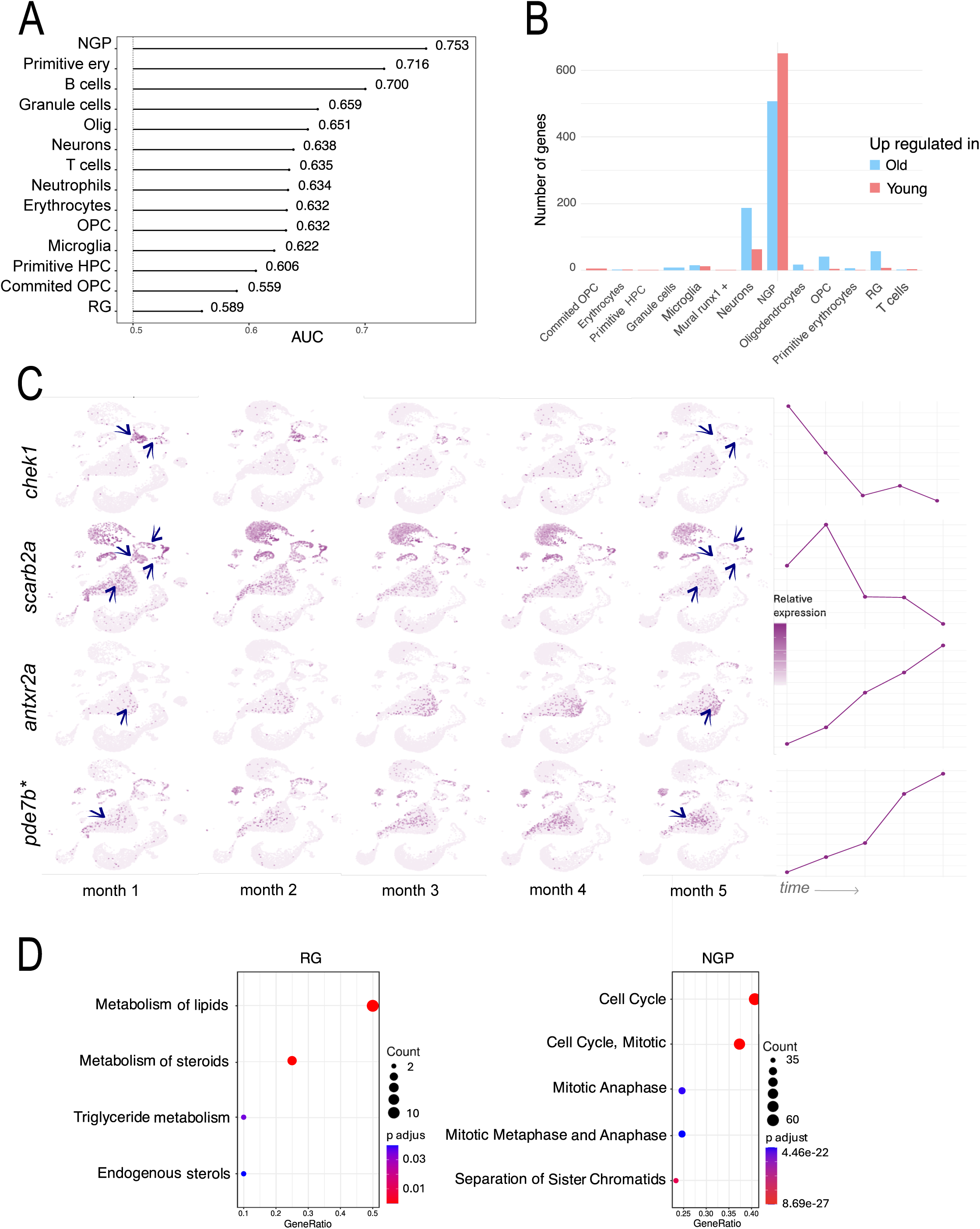
Gene expression dynamics between young and old animals. **(A)** Lollipop plot of AUGUR AUC (Area Under the Curve) results per cell type, showing the sensitivity of each cell type to the tested parameter, in this case the age of the animal. **(B)** Histogram of the number of differentially expressed genes (DEG) between old and young killifish across each major cell type. **(C)** UMAP projections separated by fish age, illustrating a progressive decrease in scarb2a and chek1 expression over time and the increase in antxr2a and pde7b expression. Arrows mark the cell clusters where the expression changes are more significant. Lineplots at the right represent the expression dynamics along the 5 timepoints.

In NGP, genes that were upregulated in old animals were enriched in GO terms related to cell proliferation (Fig. 3D and Supp. Table 4). This suggests that, in adult individuals, these cells may be exiting a state of maintenance or reserve, leading to their continuous division and eventual depletion. This aligns with the observed drastic decrease in this cell type with age. In old animals, RG exhibited enriched GO terms related to lipid metabolism (Fig. 3D and Supp. Table 4), which could be linked to oxidative stress protection due to the increased presence of reactive oxygen species (ROS) (Goodman and Bellen 2022).

### Evidences for primitive hematopoiesis in the adult killifish brain

In addition to cells of neuroectodermal origin, we also identified a significant number of clusters of hematopoietic/endothelial origin, some of which were not expected, *a priori*, to be found in the brain of adult animals. This was the case for cluster 10 (Mural runx1+) that expresses perivascular markers like *cspg4* and *acta2* together with *runx1*, a master regulator of the endothelial-hematopoietic transition (EHT) (Fig. 4A-C) (M. J. Chen et al. 2009). In mice, these cells are present in the aorta gonad-mesonephros (AGM) and are pivotal for the embryonic hematopoiesis (Gonzalez Galofre et al. 2024). More importantly, we found that some key genes expressed in cluster 10 are also expressed in cluster 3 (EC) (*Rgs5b*, *Notch1b*, *Jag1b, Acta2*). Of note, we observed that *Runx1* was differentially expressed in males and females in the EC cluster (fig. 4A, C). A closer look at cluster EC showed that not only it expresses pan-endothelial marker genes (*Pecam1/Pecam1a* and *Pecam1b*, *Cdh5, Sox17*) but also actively transcribed mesenchymal genes such as *Pdgfrb*, *Mest*, *Cd248a*, or *Acta2*. Strikingly, several genes expressed in hemangioblasts (*Runx1*, *Lmo2*, *Eomes a*, *Tal1*, *Kdrl*/*Flk1*) and hemogenic cells (*Runx1*, *Fli1*, *Notch1a*, *Jag1b*) (fig. 4A) were expressed in EC cells. These characteristics were particularly evident in the case of young females. The doublet discrimination algorithm showed there are no doublets present in this cluster, ruling out the possibility of this observation being an artifact (fig. 4B). Both in mice and humans, hemangioblasts and hemogenic cells are two different types of specialized cells that support hematopoiesis only during embryonic stages (Haniffa et al. 2024). Thus, cluster 3 and cluster 10 seem to hold “hybrid” properties of hemangioblasts and hemogenic cells. As far as we know, there is no evidence that either hemangioblasts or hemogenic cells are circulating in the bloodstream. Altogether, these data suggest that a primitive program of hematopoiesis is operating in the brain of adult killifish. Supporting this idea, we identified cluster 7, named as FGP (Fluorescent granular perithelial cells), which expresses *notch1a*, *sele* and *lmo2*. FGP are perivascular cells surrounding vessels of the blood-brain barrier and are also located in the perivascular spaces of the meninges, a place where hematopoietic stem cells are locally maintained in the brain of adult mouse (Niu et al. 2022; Brioschi et al. 2021). FGP have been also described in the brain zebrafish and previous studies have shown that they emerge by differentiation of the endothelium (Brioschi et al. 2021), raising the possibility that EC and FGP might be spatially associated. Supporting this idea, Clusters 3, 7 and 10 share the expression of several genes (fig. 4A). Of note, the hypothesis of a primitive hematopoiesis program operating in adult killifish is reinforced by the presence of cluster 6 (primitive HPC), which showed the highest score for endothelial-hematopoietic transition (EHT) and marker genes associated with GO terms related to myeloid cell differentiation (fig. 4D, E) (M. J. Chen et al. 2009) and the cluster 17 (Primitive erythrocytes) expressing hemoglobin embryonic genes (fig. 4F).

**Figure 4.**
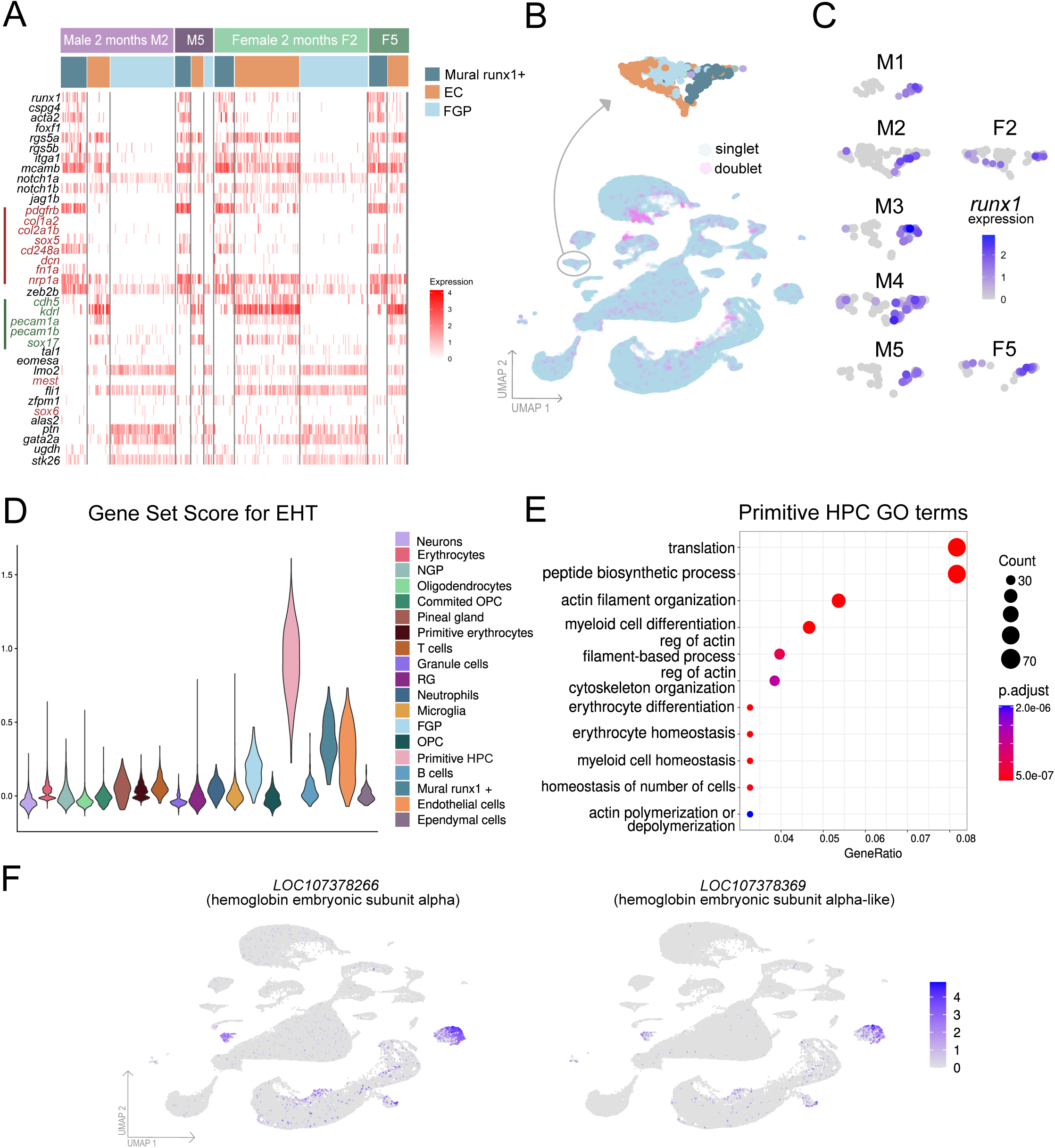
Primitive hematopoietic traits in the adult brain. **(A)** Heatmap showing the expression of genes in mural runx+ cells, EC and FGP, at 2 and 5 months in both sexes. The line in red and green indicates classic mesenchymal and endothelial marker genes, respectively. Labeled as M (male) or F (female) followed by the age in months. (**B**) UMAP displaying doublets and singlets in samples, to discard artifactual effects. (**C**) UMAP showing the expression of runx1 in runx+ cells, EC and FGP. Labeled as M (male) or F (female) followed by the age in months. (**D**) Gene set score for endothelial-to-hematopoietic transition (EHT) across cell types in the adult killifish brain. Violin plots show the distribution of scores calculated using AddModuleScore. (**E**) Dot plot showing enriched Gene Ontology (GO) terms derived from markers of Erythroid Progenitors cluster. The dot size indicates the proportion of genes associated with each term, while the color scale reflects statistical significance. (**F**) UMAP visualization depicting the expression patterns of hemoglobin embryonic subunit alpha genes LOC107378266 and LOC107378369, generated from the single-cell RNA-seq data.

As a whole, our single cell and bulk RNAseq analyses identified several groups of genes with functions related to embryonic stages, including neural development or hematopoiesis. Indeed, we found several embryonic genes that were consistently expressed in the killifish brain as it ages, which was an unexpected finding for adult animals (Supp. Fig. 5A-B). For example, the ortholog of the *crestin* gene was found to be abundantly expressed in neurons, oligodendrocytes and granule cells (Supp. Fig. A). *Crestin* is known for its role in neural crest cell migration, where it helps guide these multipotent cells to their target locations during early embryonic development (Rubinstein et al. 2000). Its expression in adult samples was unexpected, as it is predominantly associated with embryonic development rather than mature tissues. Another illustrative example of this involves embryonic hemoglobins, already mentioned previously, which are supposed to be expressed only during embryonic development (Brownlie et al. 2003; Palis et al. 2010) but appear highly expressed in females until the senescence (fig. 4D).

### Neotenic features in adult killifish transcriptional programs

We then wondered whether the expression of these embryonic genes in the adult brain could be something incidental, or represent a general trend in killifish. To evaluate this hypothesis, we performed a direct gene-to-gene comparison, using orthology conversion tables (Supp. Table 5), between our adult killifish samples and already published single cell datasets from both zebrafish adult brains and embryo heads (Raj et al. 2020) (Supp. Table 6). Strikingly, our adult killifish samples correlated better, in general, with embryonic than with adult zebrafish clusters (fig. 5A, B), which suggests that the expression of individual embryonic genes in the adult killifish brain reflects a general tendency.

**Figure 5.**
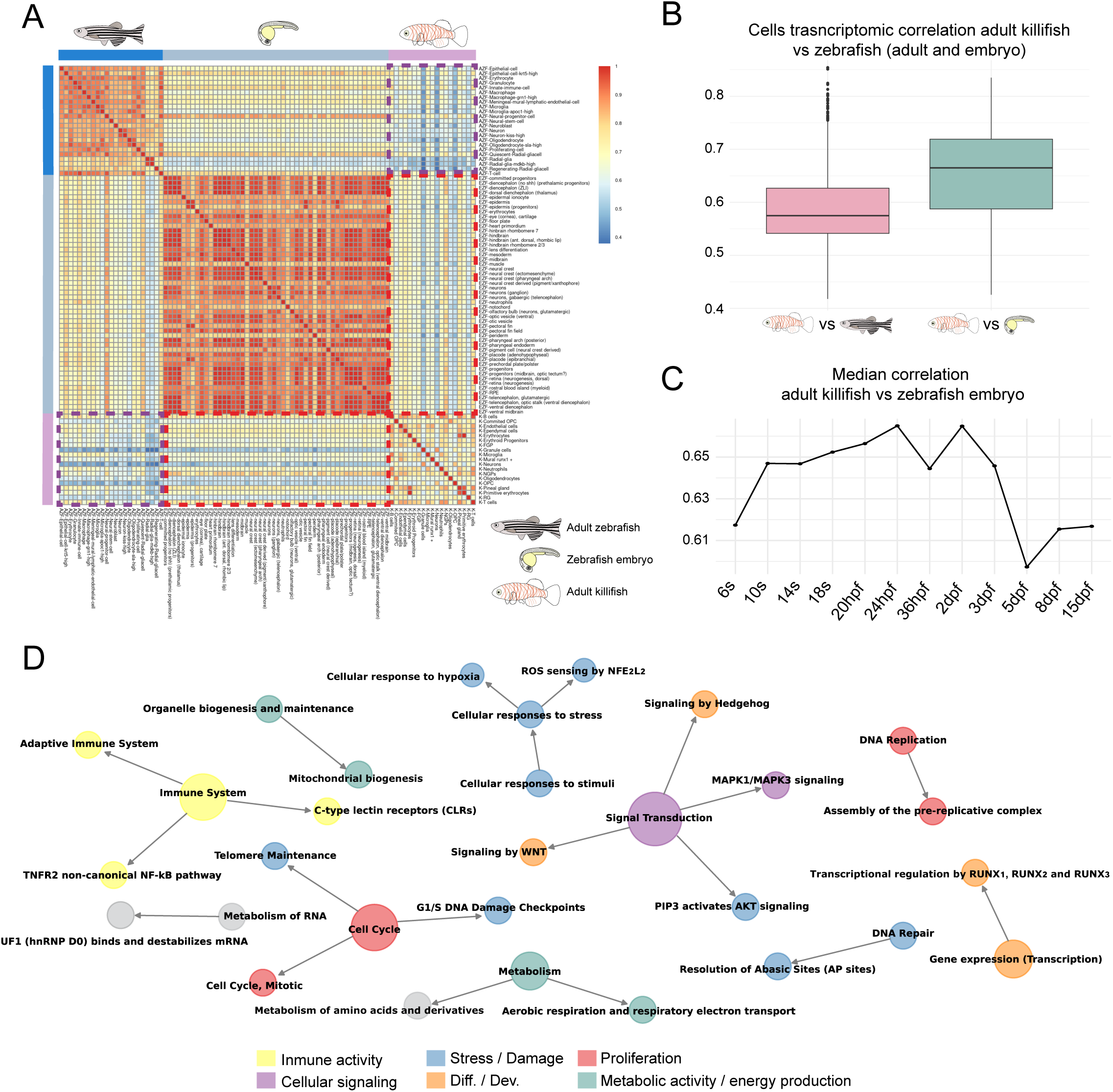
Gene expression comparison between adult killifish brain and zebrafish, adults and embryos. **(A)** Correlation in gene expression among different cell types in the brain of zebrafish embryo (ZFE), adult (AZF) and adult killifish (AKF). The region of the heatmap corresponding to the comparison of ZFE vs AKF is marked with a dashed red rectangle, and AZF vs AKF with a dashed purple rectangle. **(B)** Distribution of transcriptomic correlations between AZF vs AKF (pink) and EZF vs AKF (green) cell types. **(C)** Trend of median transcriptomic correlation between EZF and AKF over several stages of zebrafish development. **(D)** Compilation of the Reactome terms that are upregulated in both young adults and aged adults of killifish when compared to adults of zebrafish from four different strains.

To broadly assess differences in the adult killifish brain—independent of age—we analyzed whole-brain gene expression using bulk RNA sequencing and compared it to that of four adult zebrafish strains (Supp. Table 6). The differentially expressed genes (DEGs) upregulated in killifish were evaluated for enrichment using Reactome terms. This analysis consistently identified, across comparisons with all four zebrafish strains, pathways related to accelerated cell cycle and proliferation, high metabolism and energy production, and differentiation and tissue remodeling, which are typically associated with embryonic developmental programs (fig. 5D, Supp. Table 7).

## Discussion

In this study, we took advantage of scRNAseq performed in killifish brains at five different timepoints, from one month, when the animals are just sexually mature, until 5 months, when they are considered senescent. We analyzed changes in both the number of cells of different clusters and the transcriptomic level of genes in these clusters, in order to determine the cell types and the genes that are more sensitive to the aging process in the adult killifish brain. These datasets constitute an unique and rich resource to study the process of aging in the vertebrate brain.

Among the identified cell clusters, we found clusters of stromal cells (EC and Mural *runx1*+) with hybrid characteristics of hemangioblasts and hemogenic cells, which could suggest a rapid cellular transition from Mural *runx1*+ to EC as have been described in embryonic stages of zebrafish (Trinh et al. 2023). As far as we know, hemangioblasts and hemogenic cells are involved in embryonic hematopoiesis in other species and there is no evidence that these cells circulate in the bloodstream (Koeniger et al. 2021). The identification of FGP (perivascular cells) in our scRNAseq data, together with recent evidence supporting their emergence from endothelial cells (Venero Galanternik et al. 2017), suggest that these three clusters (EC, Mural *runx1*+ and FGP) could be sharing spatial localization. In addition, the high EHT score observed in cluster 6 (primitive HPC) suggests two possible scenarios: either a primitive hematopoietic program is occurring in the brain of adult killifish, similar to what happens in mouse in the AGM during embryonic stages, or EC and Mural Runx1+ cells are required to create the conditions necessary for the long-term maintenance of multipotent hematopoietic progenitors originated in the main hematopoietic tissues during development. While definitive hematopoiesis has been reported in the brain of adult mice (Koeniger et al. 2021), it was surprising to find evidence for primitive hematopoiesis in both adult and senescent killifish. The most striking finding was the identification of cluster 17 (primitive erythrocytes), which expresses embryonic hemoglobin (genes *LOC107378266* and *LOC107378369*). Interestingly, we observed that *runx1* expression in EC cluster seems to be almost exclusive to females, being hardly detected in male EC. In fact, young females displayed EC with the strongest characteristics of hemangioblasts and hemogenic cells. We wonder, whether or not, this is linked to the persistence of primitive erythrocytes in the brain of old females and is supporting local generation, in contrast to males, where this population is much less represented from the second month onwards. However, from our studies it cannot be concluded if primitive erythrocytes, as well as other hematopoietic clusters identified, emerge from brain hematopoiesis or are coming from the bloodstream.

The presence of primitive erythrocytes in the brain of female killifish could confer an advantage, as embryonic hemoglobin has higher oxygen affinity than adult hemoglobin, playing a neuroprotective role against oxidative stress resulting from the high metabolic rate of the brain. In fact, female killifish reach the senescence in better physical conditions than males (Di Cicco et al. 2011). A non-exclusive hypothesis would be that high oxygen affinity represents an adaptive mechanism to low O2 concentration in the warm and stagnant waters where killifish live. Furthermore, primitive progenitors can generate a wide variety of blood cells, specially under stress conditions, which could enhance the immediate immune response to rapidly respond to infections or injuries.

Future studies, focused specifically on brain hematopoiesis, will be required to establish whether primitive hematopoiesis is occuring in the brain of adult killifish and what type of hematopoietic cells are being generated.

Concerning the neural compartment, our cell-fate analysis predicts that NGP give rise not only to RG cells and neurons, as was previously reported, but also to OPC, responsible of the generation of oligodendrocytes, which are essential cells that generate myelin for the insulation of axons in the central nervous system, during the adulthood. This could be a specific phenomenon in killifish, since OPC are produced from different progenitor cells only during embryonic development in other species (Marques et al. 2018).

Single cell RNAseq results were supported by bulk RNAseq experiments in the same tissue. We found several synexpression groups of genes that confirm the results obtained from our single cell experiments and from previous assays from other laboratories (Baumgart et al. 2014). For example, we identified a group of genes with high expression at 1.5 months that decreases at 2.5 and 4 months, but increases again at 5 months (group 14). This cluster was enriched in GO terms related to biosynthesis of proteins, which is high in young animals and lower in more mature individuals. However, it has been previously reported that the efficiency of protein synthesis is much lower in aged animals, and this is compensated by an important increment in the synthesis rate, which produces a loss of proteostasis during senescence (Kelmer Sacramento et al. 2020; Munkácsy et al. 2019; Rodriguez, Gaczynska, and Osmulski 2010). The gene expression profile observed in the mentioned gene cluster reflects this impaired proteostasis.

The gene group 16 was composed of genes with high expression levels at the two first stages and was enriched in GO terms related to cell division. This could correspond to progenitor cells that are more active in young animals, and their activity and number are decaying during adulthood.

Finally, the gene group 1, with high expression only at the first time point was enriched in GO terms compatible with early development. Despite this being our first time point and it is related with the onset of sexual maturity, the expression of these genes involved in early development should be extinguished by this time.

Overall, our findings are compatible with the notion that the killifish brain displays neotenic features at the gene expression level for a more prolonged period, and even being maintained for some programs during the whole life span. This hypothesis was supported by the direct comparison of our single cell clusters with embryonic and adult zebrafish samples, in which we could observe a higher correlation with the former than with the latter. Furthermore, comparing bulk RNAseq from adult killifish and zebrafish highlighted biological functions related with early development, proliferation and differentiation as the most prevalent in killifish compared to zebrafish.

In all vertebrate species, the embryonic period is characterized by a high cell proliferation rate compared to the adult phase (Ruiz et al. 2011). Besides, in some species like salamanders and frogs, increasing the length of the larval period leads to increased body size (Bonett et al. 2022; Bruce 2013; Phung et al. 2020). This supports the hypothesis that the observed neotenic status in the adult killifish brain might be a necessary adaptation to sustain a very high level of metabolism and cell proliferation. However, it is also known that embryonic genes are reactivated in adult tissues to respond to an injury, in the posterior process of reparation and regeneration (Albano and Hackam 2024; Poplawski et al. 2020; Zheng and Tuszynski 2023). Thus, the expression of embryonic genes in the adult killifish brain could also help mitigate the challenges associated with its high metabolic rate and its potential consequences. Whether the transcriptomic neoteny reported in this study reflects the first or the second case, or both, cannot be inferred from our results.

Our analysis helps to understand the evolutionary mechanisms that cells need to acquire in order to adapt to the species’ habitat, and suggests that, although killifish is a very good model to investigate aging and age-related diseases, extrapolation of results obtained in this species to other vertebrate species, like mammals, should be carefully addressed. However, the neotenic characteristics observed in the adult killifish brain might be part of the species’ adaptation to its demanding ecological niche, and could help identify promising cellular/molecular targets for regenerative or anti-ageing therapies in the near future.

## Materials and Methods

### Animal experimentation

All experiments involving *Nothobranchius furzeri* performed in this work conformed European Community standards for the use of animals in experimentation and were approved by the Ethical Committees from the Universidad Pablo de Olavide, Consejo Superior de Investigaciones Científicas (CSIC) and the Andalusian Government under the Ethics Agreement 16/10/2020/007.

### Fish stocks

Turquoise killifish of the MZM-0410 strain were raised, maintained and bred under standard conditions (Dodzian et al. 2018; J. Chen, Khondker, and Brunet 2023). Water parameters and feeding protocols were kept strictly constant among different batches of fish to ensure that differences among adults were due to individual intrinsic variations.

### Quantitation of Transcript Levels by RT-qPCR

Adult fish were raised until they were sacrificed at different temporal points. At these points, fish were deeply anesthetized with a solution containing 168 mg/l of Tricaine (Ethyl 3-aminobenzoate methanesulfonate salt, Merck, ref. MS-222) and afterwards they were transferred to a solution containing a higher concentration (1.5 g/l) to sacrifice them (Lee and Kim 2021).

Brains were dissected and individually homogenized in TRIsure (Bioline, ref. BIO-38032) to extract total RNA with Direct-zol RNA MiniPrep kit (Zymo Research, ref. R2052) according to the manufacturer’s protocol.

cDNA was synthetized using High Capacity cDNA Reverse Transcript kit (Applied biosystems-Thermo Fisher Scientific, ref. 4368814). The expression levels of genes of interest in killifish brains were quantified through RT-qPCR (CFX96 real-time C1000 thermal cycler, Bio-Rad) and normalized to the expression level of the housekeeping gene ef1a (see primers at Supp. Table 8). qPCR reactions were performed in triplicate using iTaq Universal SYBR green Supermix (Bio-Rad, ref. 172-5124) following the manufacturer’s instructions. The expression levels for these genes in aged brain samples were calculated in relation to young ones (average set to 100%).

### scRNAseq sample preparation

Killifish were raised until desired age and sacrificed as described above. For each sample, two adult brains were dissected and processed for single cell preparation as described in the “Adult Brain Dissociation Kit for mouse and rat” protocol (Miltenyi Biotec, 130-107-677). For our samples, we ran the “37C_ABDK_02” program and worked with volumes corresponding to the option of 20-100 mg of starting material during the debris removal step. The red blood cell removal step was skipped.

After cleaning the cells from debris, they were pelleted and resuspended in 1 ml of HBSS – Fetal Bovine Serum (FBS) 2% – Hepes 10 mM pH 7.4 – 1X Penicillin/Streptomycin (HBSS, Gibco, ref. 14175095; FBS, Gibco, ref. A3840401; Hepes, Gibco, ref 15630049; Penicillin-Streptomycin, Gibco, ref. 15140122). Cells were filtered using a 45 µm cell strainer and 500 µl of them were FACS-sorted with a SONY MA 900, using a 100 mm nozzle. Low sample pressure (20.10-20.30 psi) and the software SONY-MA were used for designing gate strategy. Dead cells were excluded by using the viability dye 7AAD (7-aminoactinomycin D, Biolegend ref. 420404) using the laser 488 and the FL4 channel (695/50). To enrich the sample in less frequent cell populations, two gates based on FFC_A and SSC_A were designed, one for big, less frequent cells and another for smaller, more abundant cells. Cells were collected in polypropylene tubes previously coated with HBSS – FBS 10%,Hepes 10 mM pH 7.4 – 1X P/S. From 8-10 times more cells than required for scRNAseq processing were sorted. Reanalysis was performed in the two types of sorted cells with purity >90%. These cells were mixed in equal proportion. The same conditions and gate strategy were maintained for the sorting of all samples prepared on different days. Cell viability, cell aggregates and counting were assessed in a TC-20 BioRad Cell Counter. Viability of cell samples was around 80%.

After sorting, cells were centrifuged at 4°C, 500 g, 7 min and gently resuspended in HBSS – FBS 2% – Hepes 10 mM pH 7.4 – 1X P/S. Then, they were filtered using a Flowmi (SP Bel-Art) filter and finally diluted at a concentration of 800-1200 cells/µl. A total number of 20.000 cells were loaded in a Chromium 10X Genomics.

The scRNAseq libraries were constructed using Chromium Single Cell 3’ v3 Reagent Kit according to the manufacturer’s protocol. Amplified cDNA and final libraries were analyzed in a Bioanalyzer 2100 with the High Sensitivity DNA kit (Agilent Technologies). Libraries were sequenced on a DNBSEQ-T7 system (PE100/100/10/10).

### Data Processing, Quality Control and Filtering

The single-cell RNA sequencing (scRNA-seq) data were processed using **Cell Ranger v7.2.0** with the genome assembly **UI_Nfuz_MZM_1.0**. To generate a custom reference genome, the cellranger mkgtf function was used to filter the original genome annotation file. This step retained only protein-coding genes and long non-coding RNAs (lncRNAs) by specifying the relevant attributes. The filtered annotation, together with the genome sequence, was then used to build a Cell Ranger-compatible reference genome using the cellranger mkref function. This custom reference genome enabled accurate mapping and quantification of transcriptomic reads.

Raw FASTQ files for seven biological samples were processed using the cellranger count function, which aligns sequencing reads to the reference genome, quantifies gene expression, and filters out low-quality barcodes. Sample-specific gene-cell matrices were generated, and all matrices were merged to create a combined dataset for quality control and downstream analyses.

Quality control was performed using **Seurat v5.0.2** (Hao et al. 2024) to ensure the inclusion of high-quality cells and biologically relevant genes. A Seurat object was created with the CreateSeuratObject function, retaining only genes expressed in at least 10 cells across the dataset. Cells were filtered using the subset function based on stringent criteria: cells with fewer than 400 detected genes, more than 15% mitochondrial reads, or total read counts outside the range of 800 to 60,000 were excluded. Additionally, clusters containing fewer than 30 cells across all seven samples were removed to ensure robust representation of cell populations.

### scRNAseq primary analysis

Normalization of the data was performed using the SCTransform function in Seurat. This method corrects for technical variability, including differences in mitochondrial read content, by regressing out these effects. The resulting normalized dataset was used for dimensionality reduction, clustering, and visualization.

To address potential batch effects among the seven biological samples, data integration was performed using **Harmony v1.2.0** (Korsunsky et al. 2019). The RunHarmony function was applied to align datasets from all samples into a shared low-dimensional space. The integrated dataset was then subjected to dimensionality reduction, clustering, and visualization using the RunUMAP, FindNeighbors, and FindClusters Seurat functions. A clustering resolution of 0.55 was used to delineate cell populations in the integrated dataset, ensuring consistency across samples.

Cell type annotation was carried out using known marker genes specific to distinct cell types: *gja1b* for RG, *elavl4* for neurons, *gata1a* for erythrocytes, *p2ry12* for microglia, *mpa* for oligodendrocytes, *neurog1* for NGP, *cd4-1* for T cells, *LOC107378266* for primitive erythrocytes, *asmt* for pineal gland cells, *sema5a* for OPC, *mpx* for neutrophils, *acta2* for mural cells, *zic1* for granule cells, *mrc1a* for FGP, *sox10* for committed OPCs, *epd* for ependymal cells, *kdrl* for endothelial cells, and *cd79a* for B cells.

After completing cell type annotation, marker genes were identified for each population using the Seurat **FindAllMarkers** function with only.pos = TRUE (Supp. Table 3).

To calculate the endothelial-to-hematopoietic transition (EHT) score, genes associated with endothelial (*pecam1a*, *cldn5b*, *esama*, *f13a1b*), hematopoietic (*runx1*, *gata2b*, *tal1*, *mpl*), transitional (*fli1*, *tgfb1a*, *ccn2a*, *acta2*), migration (*tagln*, *cald1a*, *vclb*, *rhof*) and vascular (*zfpm1*, *f2rl2*, *itgb1a*) characteristics were selected. The AddModuleScore function in Seurat was used to integrate the average expression of these genes across cells, generating a score reflecting their association with the EHT process. This score was then used for comparative analyses across cell types.

### Cell Population Dynamic Analysis

For cell population dynamic analysis, we used **scProportionTest** (version 0.0.0.9) (Miller et al. 2021), **scVelo** (version 0.3.1) (Bergen et al. 2020), and **CellPhoneDB** (version 5.0.1) (Vento-Tormo et al. 2018). First, we subset the dataset by sex, focusing on female and male populations, and performed pairwise comparisons between the age groups “month_2” and “month_5” using permutation tests with 100 bootstraps, applying significance thresholds of FDR = 0.025 and fold change = 2. For cell fate dynamics, **scVelo** was applied to a subset of cell types, including NGP, OPC, and RG, starting with normalization and filtering via pp.filter_and_normalize function, followed by stochastic velocity estimation and velocity graph computation. Additionally, **CellPhoneDB** was employed to analyze cell-cell interactions using human orthologs (GRCh38.p14), with results visualized via **ktplots** (version 2.3.0).

### RNA-seq experiments

Brain tissue was prepared as described above for RT-qPCRs. Forebrain, midbrain and hindbrain from males were dissected and processed individually. Tissue samples were dissected and homogenized in TRIsure (Bioline) and RNA was extracted and purified with Direct-zol RNA Miniprep kits (ZYMO). Total RNA was sent for library preparation and sequencing.

### Pseudo-bulk and bulk-RNA-seq analysis

For the pseudobulk RNA-seq analysis, data from single-cell RNA-seq experiments were processed using the muscat package (version 1.8.2). The dataset included single-cell profiles from four samples: two males aged 2 and 5 months and two females aged 2 and 5 months. Quality control was applied to exclude cells with extreme gene detection levels and lowly expressed genes. Differential expression analysis was conducted with the pbDS function using a contrast matrix to compare age groups. Differentially expressed genes (adjusted p-value < 0.05, log fold change > 0.9) for each cell type across time were subjected to Gene Ontology (GO) and Reactome pathway enrichment analyses using zebrafish orthologs with clusterProfiler (version 4.2.2) (Yu et al. 2012) and ReactomePA (version 1.38.0) (Yu and He 2016).

For the bulk RNA-seq analysis, raw sequencing reads were aligned to the *Nothobranchius furzeri* genome assembly (UI_Nfuz_MZM_1.0) using STAR (version 2.7.10a) (Dobin et al. 2013), followed by quantification of gene expression with featureCounts (version 2.0.6) (Liao, Smyth, and Shi 2014). Quality control ensured the exclusion of low-quality samples. Data from four time points—8, 12, 20, and 24 weeks—were analyzed using DESeq2 (version 1.34.0) (Love, Huber, and Anders 2014), with each time point compared to the 8-week baseline using a design matrix. Genes showing significant changes (adjusted p-value < 0.05, absolute log2 fold change > 0.4) were clustered into 20 groups using k-means clustering, revealing dynamic temporal expression patterns. Enrichment analyses for GO terms and Reactome pathways were performed with clusterProfiler (version 4.2.2) and ReactomePA (version 1.38.0) using zebrafish orthologs.

### Comparative analysis between zebrafish and killifish

To assess the transcriptional similarity between adult killifish, adult zebrafish, and embryonic zebrafish cell types, we analyzed single-cell RNA sequencing (scRNA-seq) data from publicly available datasets. Gene nomenclature discrepancies were addressed by mapping killifish gene names to zebrafish orthologs generated using Orthofinder (Emms and Kelly 2019). We performed data normalization, integration, and dimensionality reduction using Seurat, followed by correlation analysis between cell type expression profiles.

To identify Reactome terms upregulated in the brain of killifish compared to zebrafish, we performed differential expression analysis (DEA) using DESeq2 (version 1.34.0) in R across multiple comparisons. Four different strains of zebrafish at 17 weeks old (GSE61108) were compared to killifish samples at different stages (8 and 24 weeks). Raw read counts were processed and normalized, and genes with low expression (total counts <10) were filtered out. Orthologous genes between species were identified using OrthoFinder. Significantly differentially expressed genes (DEGs) were identified based on p.adjust < 0.05, followed by functional enrichment analysis using clusterProfiler (version 4.2.2). Significantly enriched Reactome terms were extracted from the DEGs, and to control for age-related bias in killifish, we focused on Reactome pathways consistently upregulated at both 8 and 24 weeks, identifying biological processes representative of the general adult stage.

## Acknowledgments

This work was supported by the Spanish Ministerio de Ciencia e Innovación (PID2022-141288NB-I00 to JJT and PID2023-148267NB-I00 to JLR) and CEX2020-00108-M Unidad de Excelencia María de Maeztu.

## Data availability

Original data presented in this study have been deposited in GEO Datasets repository, under accession numbers GSE290929 and GSE290930.

## Author contributions

AMB analyzed the data and interpreted the results with the help of JT, SN, GN and JLR. SN, LA, VH, MMO, ARB and AFM performed the scRNAseq, RNAseq and RT-qPCR experiments. JT conceived the study and wrote the manuscript with the help of GN, JLR, SN and AMB.

## Conflict of interests

The authors declare no competing interests.

## Notes

### Competing Interest Statement

The authors have declared no competing interest.

