## Supplementary Figures for "Neotenic transcriptomic features in the adult turquoise killifish brain"

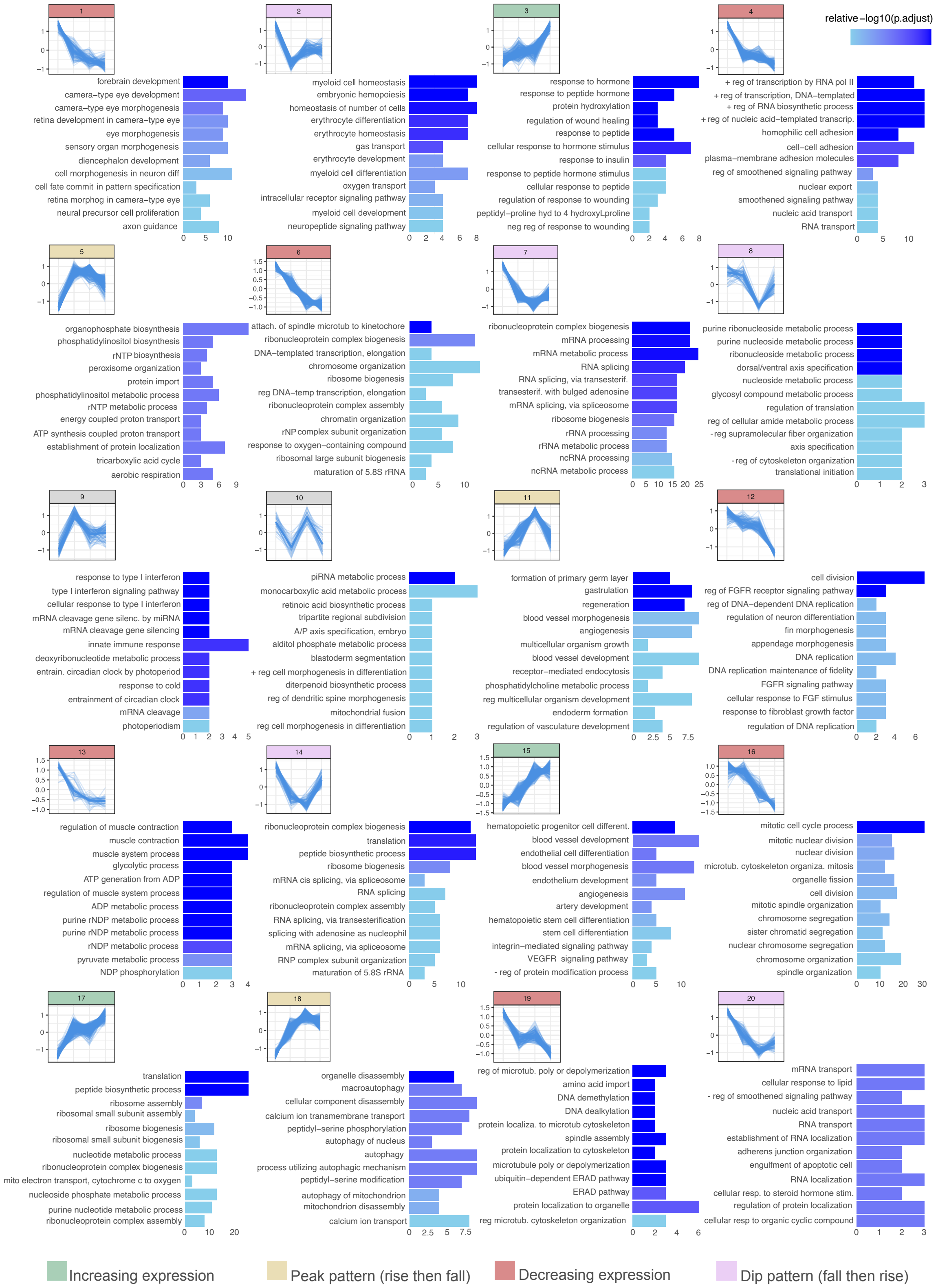


**Supplementary Figure 1. Gene synexpression groups from RNAseq analysis.** Differentially expressed genes (DEGs) identified through RNA-seq analysis of four timepoints, comparing each time point to the baseline. Temporal expression dynamics were grouped into 20 groups using k-means clustering. Each panel shows the average expression trend of genes over the four timepoints (1.5, 2.5, 4 and 5 months) in each cluster, along with the top 12 associated Gene Ontology (GO) terms (Biological Process).

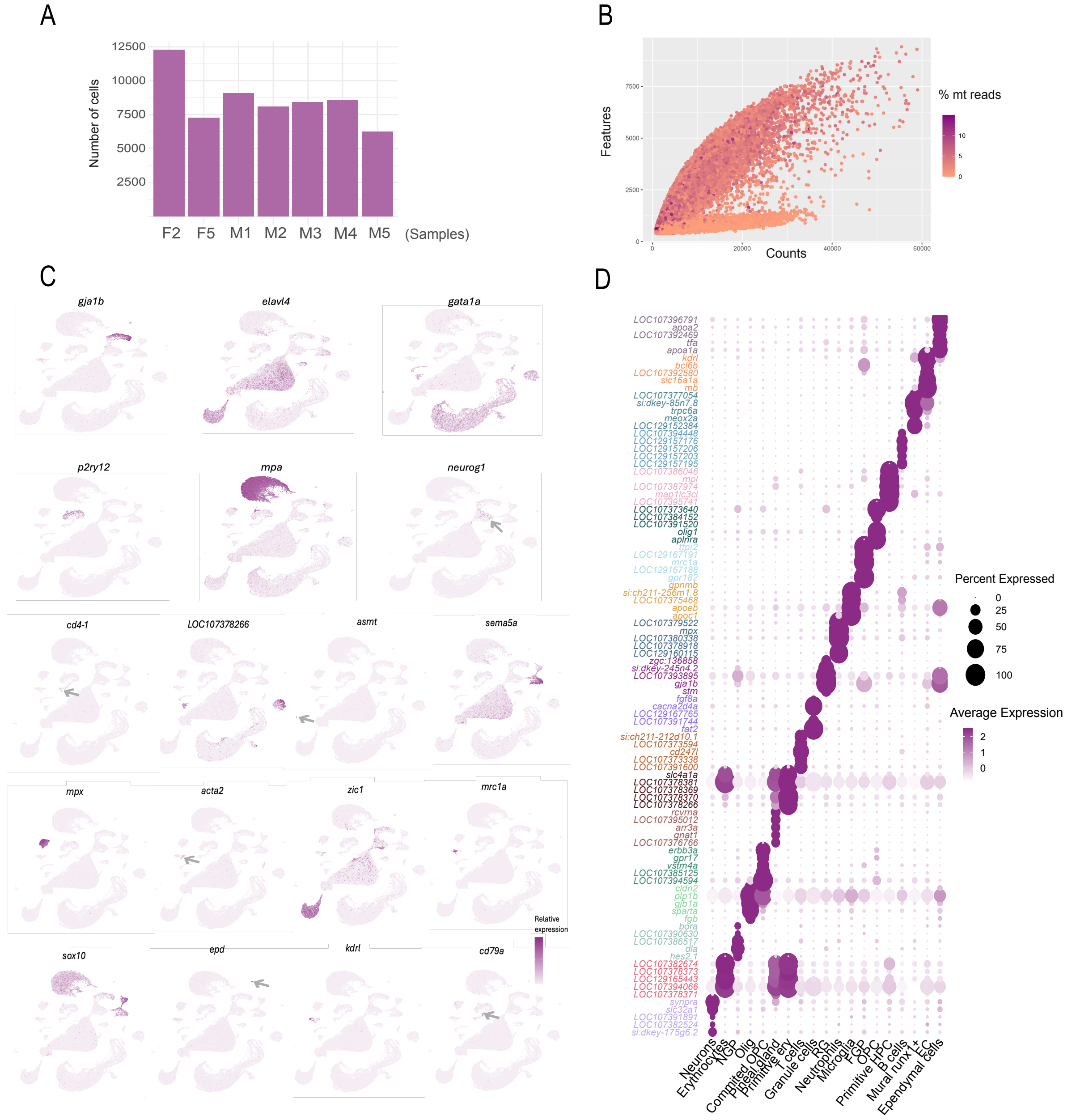


**Supplementary Figure 2. scRNAseq metrics, annotation and marker genes. (A)** Histogram showing the distribution of cells across the seven samples. The y-axis represents the number of cells, while the x-axis corresponds to the different samples, labeled as M (male) or F (female) followed by the age in months. **(B)** Scatter plot summarizing the post-filtering quality control (QC) metrics for single cells across all samples. Each point represents a cell, with the x-axis indicating the number of counts, the y-axis the number of genes, and the color scale representing the percentage of mitochondrial reads per cell. **(C)** UMAPs with the expression patterns of key marker genes used for cell type annotation, ordered as follows: *gja1b* for RG, *elavl4* for neurons, *gata1a* for erythrocytes, *p2ry12* for microglia, *mpa* for oligodendrocytes, *neurog1* for NGP, *cd4-1* for T cells, *LOC107378266* for primitive erythrocytes, *asmt* for pineal gland cells, *sema5a* for OPC, *mpx* for neutrophils, *acta2* for mural cells, *zic1* for granule cells, *mrc1a* for FGP, *sox10* for committed OPC, *epd* for ependymal cells, *kdrl* for endothelial cells, and *cd79a* for B cells. **(D)** Dot plot summarizing the expression of selected marker genes across the major cell types identified in the killifish brain. The size of each dot represents the proportion of cells within a given cell type expressing the marker gene, while the color intensity reflects the average expression level.

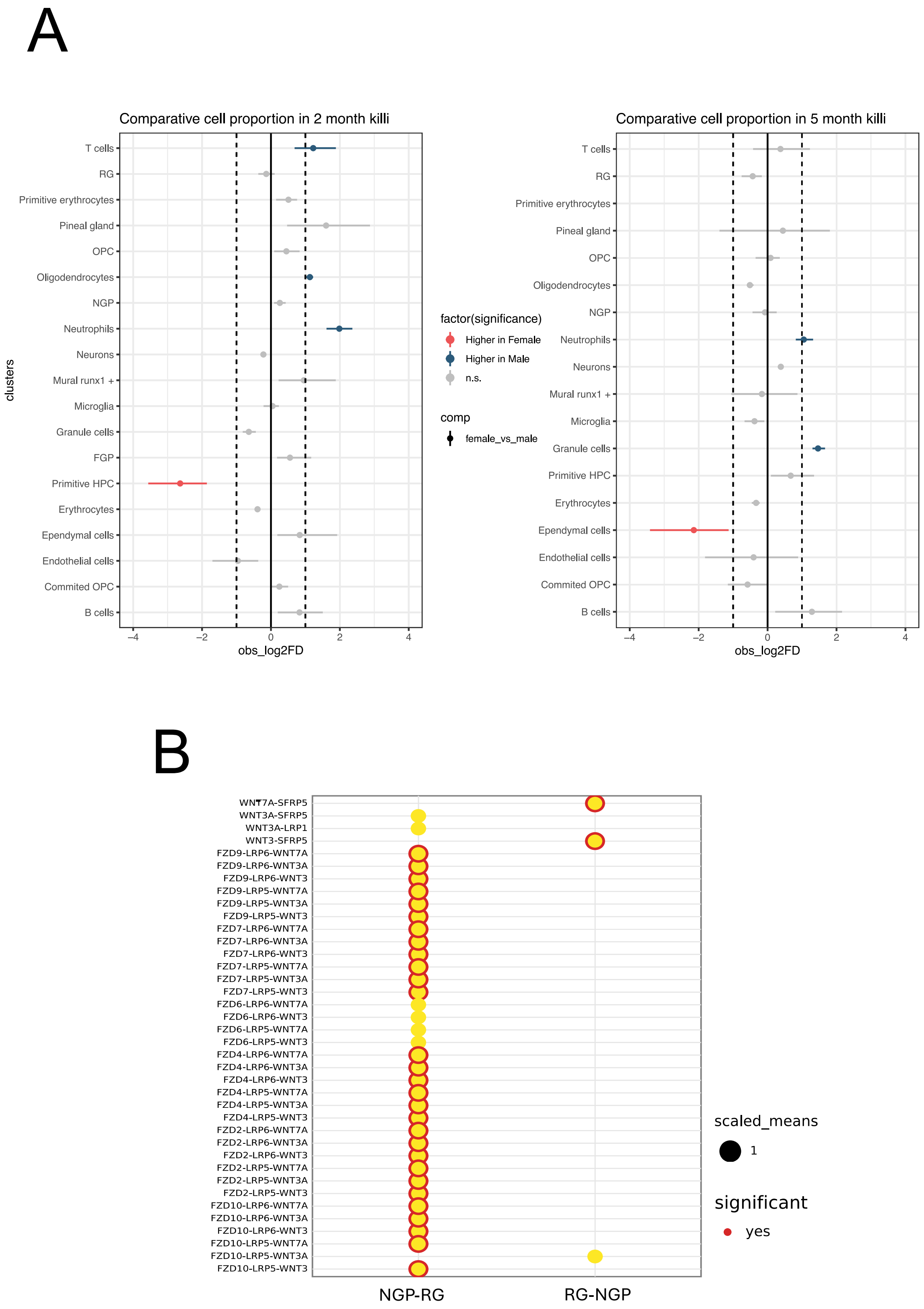


**Supplementary Figure 3. Comparative of cell population sizes. (A)** Differences in cellular population sizes between males and females at 2 months (left) and 5 months (right). Dashed lines represent significance thresholds of a 1.5-fold change in cell abundance at $p < 0.05$. **(B)** Wnt signaling pathway-related ligand-receptor interactions identified between radial glia (RG) and neural progenitor cells (NGP). The y-axis represents different ligand-receptor combinations, with red-outlined dots indicating statistically significant interactions ($p < 0.05$).

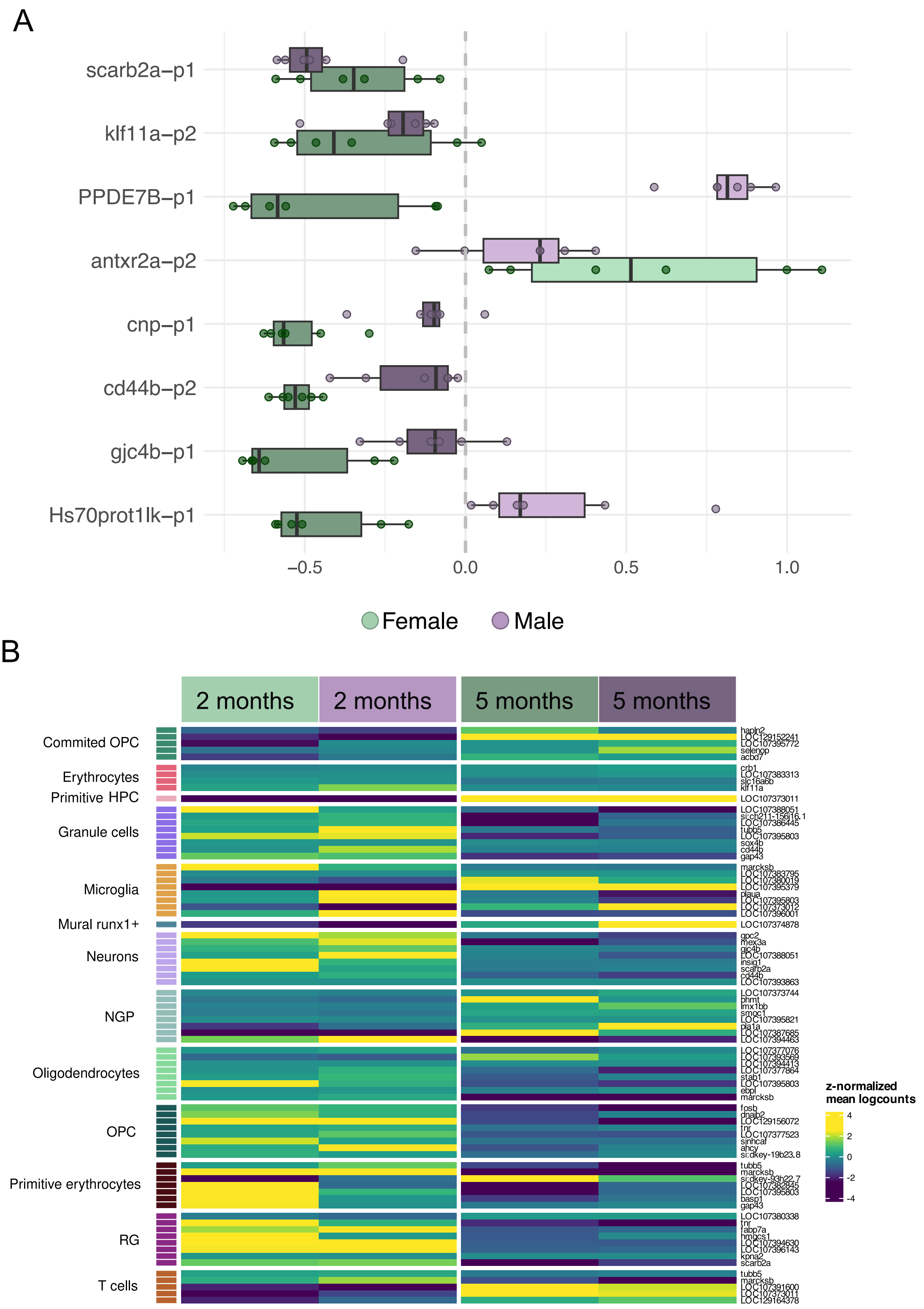


**Supplementary Figure 4. (A)** qPCR results using selected genes with changing expression profiles in scRNAseq experiments. Young (2 months) and old (5 months) females (in green) and males (in purple) have been compared. The ratio of the expression at 5 months related to the expression at 2 months is represented, so negative values represent higher expression in young animals, and positive values represent higher expression in old animals. **(B)** Top 8 DEG between old (5 months) and young (2 months) adult killifish calculated using pseudo bulk-RNAseq with muscat. Some cell types have less than 8 DEG.

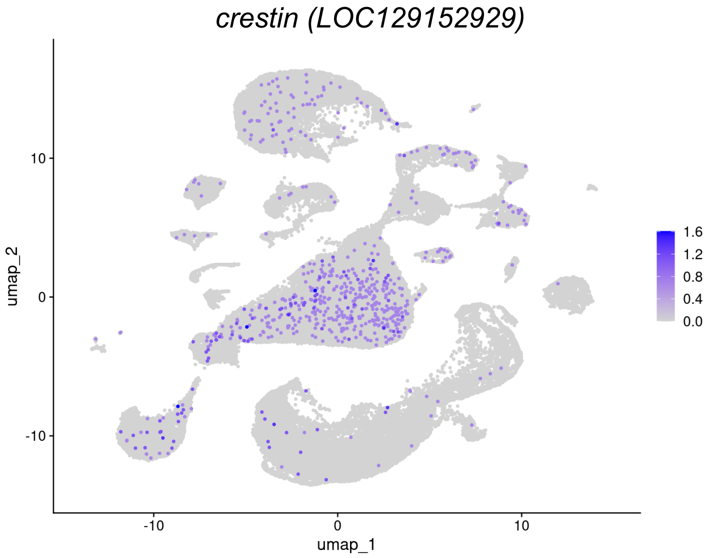


| | |
|---|---|
| *hoxb1a, hoxc6b, hoxd4a, pax2b, pax7a, pax7b* | Regulate neural progenitor fate and brain regionalization. |
| *neurod1, neurod2, dcx, neurod4, neurod6, aspm* | Control neuron differentiation and migration. |
| *svopl* | Involved in synapse formation and neural connectivity. |
| *fgf10a, fgf24* | Promote neuronal survival, growth, and differentiation. |
| *crestin, sox10* | Essential for neural crest-derived neurons and glia. |
| *chrnd, chrng* | Components of cholinergic synapses in developing neurons. |
| *dnmt3ba, dnmt3bb.2* | Establish DNA methylation patterns in neural cells. |

**Supplementary Figure 5. (A)** UMAP with the expression of *crestin* (*LOC129152929*) in the scRNA-seq samples. **(B)** Developmental genes found in adult killifish scRNAseq and bulk RNAseq using zebrafish orthologs, and their main known functions.
